# Information Flow in the Fibroblast Growth Factor Receptor Communication Channel

**DOI:** 10.1101/2021.08.23.457435

**Authors:** José Díaz, Gustavo Martínez-Mekler

**Affiliations:** Laboratorio de Dinámica de Redes Genéticas, Centro de Investigación en Dinámica Celular, Universidad Autónoma del Estado de Morelos, Cuernavaca, Morelos, México; Instituto de Ciencias Físicas, Universidad Nacional Autónoma de México, Campus Cuernavaca, Cuernavaca, Morelos, México; Centro de Ciencias de la Complejidad, Universidad Nacional Autónoma de México, Ciudad de México, México

**Keywords:** MAPK, C-myc, Fibroblast Growth Factor, Nonlinear dynamics, Information Theory, Gene Regulatory Networks

## Abstract

In this work we analyze the flow of information through the Fibroblast Growth Factor Receptor (FGFR) communication channel when different types of signals are transmitted by the MAPK cascade to the gene regulatory network (GRN) formed by the genes *C*-*Myc, DUSP*, and *Cdc25A*, which control fibroblast proliferation. We used the canonical mathematical model of the MAPK cascade coupled to a stochastic model for the activation of the gene regulatory network, subject to different types of FGF inputs (step, quadratic pulses, Dirac delta, and white noise), in order to analyze the response of the gene regulatory network to each type of signal, and determine the temporal variation of the value of its Shannon entropy in each case. Our model suggests that the sustained activation of the FGFR communication channel with a step of FGF > 1 nM is required for cell cycle progression and that during the G1/S transition the amount of uncertainty of the GRN remains at a steady value of ∼ 2.75 bits, indicating that while the fibroblast stimulation with FGF continues the G1/S transition does not require an additional interchange of information between the emitter and the gene regulatory network to be completed. We also found that either low frequency pulses of FGF or low frequency noise, both with a frequency *f* ≤ 2.77 Hz, are not filtered by the MAPK cascade and can modify the output of the communication channel, i.e., the amount of the effector proteins c-myc, cdc25A and DUSP. An additional effect suggested by our model is that o low frequency periodic signals and noise possibly blockage cell cycle progression because the threshold value concentration of cdc25A for the G1/S transition is not sustained in the in the nucleus during the 10 hours that this process lasts. Finally, from our model we can estimate the capacity of this communication channel in 0.96 bits/min.

## 1 Introduction

There are three subgroups of the Fibroblast Growth Factor (FGF) family: the canonical FGFs, the intracellular FGFs and the hormonal FGFs. The canonical FGF comprises about 20 isoforms; they exert their action by short range diffusion in the extracellular medium, and can act in a dose-dependent fashion (Böttcher, and Niehrs, 2005). Canonical FGFs are secreted ligands that bind to their specific cell surface receptor belonging to the tyrosine kinase family of FGF receptors (FGFRs). FGFs bind to their receptors with different affinity (*k*_*d*_) values that ranges from 6.19 × 10^−8^ M to 1.65 × 10^−7^ M (Mohammadi et al., 2005; Gillespie et al, 1989). The four FGFR isoforms are very similar in their global structure with an extracellular-ligand-binding domain, a single transmembrane domain and an intercellular tyrosine kinase domain (Figure 1a). The extracellular part of the receptor has two or three immunoglobulin (Ig)-like domains, and a heparin-binding domain which is important for the interaction with the ligand. This extracellular domain suffers multiple alternative splicing, which modulate the affinity of the receptor to its ligand. Between the extracellular domains D1 and D2 (Figure 1a) there is a consecutive file of acidic amino acids building up the acidic box, which is an exclusive feature of these receptors (Böttcher and Niehrs, 2005; Mohammadi, et al., 2005). The acidic box is connected to the ligand binding portion of the receptor, which consists of two loops formed by domains D2 and D3. The heparin-binding site is located in de D2 loop. The tyrosine kinase domain of the receptor is located in the cytoplasm (Figure 1a).

**Figure 1.**
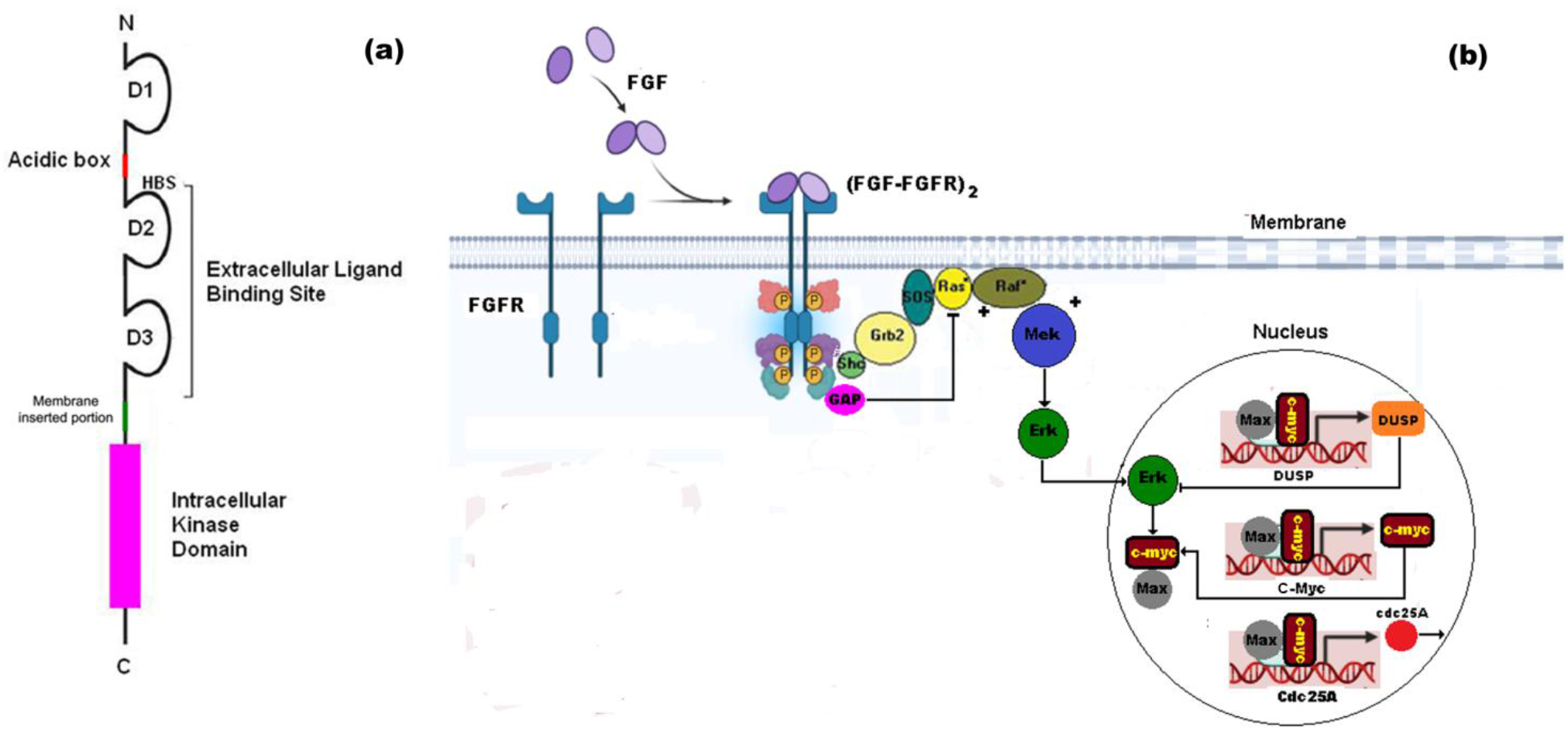
FGFR communication channel. a) The four FGFR isoforms are very similar in their global structure with an extracellular-ligand-binding domain, a single transmembranal domain and an intercellular tyrosine kinase domain. b) After the cross-phosphorylation of the FGF-FGFR complexes, the chemical signal generated by the external FGF concentration is transmitted to the small G-protein Ras through the docking proteins Shc, Grb2 and SOS. The activation of Ras is regulated by GAP, which is activated by the (FGF-FGFR)_2_ complex. Ras transmits the signal emitted by dimeric FGFR downstream through a MAPK cascade in which the MAPKKK (MAP-Kinase-Kinase-Kinase) is Raf1. Activated Raf specifically phosphorylates either MEK1 or MEK2, which is a MAPKK (MAP-Kinase-Kinase). MEK phosphorylates tyrosine and threonine residues of a MAPK (MAP-Kinase), which can be either ERK1 or ERK2; a fraction of ERK is translocated into the nucleus and activates genes such as *C*-*Myc, DUSP*, and *Cdc25A* that control the cell cycle. This series of coupled chemical processes form the FGFR communication channel in which FGF is the input signal, the (FGF-FGFR)_2_ dimer is the emitter and encoder, the MAPK cascade is the transmitter (SN), the GRN is the receiver and decoder, and the new synthesized proteins are the effectors or output.

When FGF binds to its specific receptor (FGFR) with tyrosine-kinase activity, the dimeric complex (FGF-FGFR)_2_ is formed. Figure 1b shows that (FGF-FGFR)_2_ phosphorylates the docking Sh2-Sh3 adaptor protein Shc, which binds to the phosphorylation sites Y653 and Y654 of the dimeric complex. Also, (FGF-FGFR)_2_ phosphorylates GTP-ase Activating Protein (GAP), which binds to the phosphorylation sites Y653 or Y654 of the dimeric complex (Klint, and Claesson-Welsh, 1999). After the cross-phosphorylation of the FGF-FGFR complexes, the chemical signal generated by the external FGF concentration is transmitted to the small G-protein Ras through the docking proteins Shc, Grb2 and SOS (Figure 1b). The activated state of Ras is regulated by GAP, which is activated by the (FGF-FGFR)_2_ complex. Once Ras is activated, the signal is transmitted downstream through the MAPK cascade in which the MAPKKK (MAP-Kinase-Kinase-Kinase) is Raf1. This serine/threonine kinase is translocated to the inner side of the cell membrane where it is activated by Ras, in presence of phosphatidic acid. Activated Raf specifically phosphorylates either MEK1 or MEK2, which is a MAPKK (MAP-Kinase-Kinase). MEK phosphorylates tyrosine and threonine residues of a MAPK (MAP-Kinase), which can be either ERK1 or ERK2; a fraction of ERK is translocated into the nucleus and activates genes such as *C*-*Myc* (Garrington and Johnson,1999; Arkun and Yasemi, 2018), *DUSP* (Chappell et al., 2013) and *Cdc25A* (Dang, 1999).

*C-Myc* gene belongs to the family of *Myc* genes that includes *B-Myc, L-Myc, N-Myc*, and *S-myc. C-Myc* has a central role in the regulation of cell proliferation. This gene is located on chromosome 8 in humans and is composed by three exons. The transcription factor c-myc, product of *C-Myc*, has a transactivation domain within its first N-terminal 143 amino acids, and two canonical DNA binding sites or E-boxes (5’-CACGTG-3’). Serine 62 of protein c-myc is phosphorylated by ERK to form a heterodimer with Max protein in the fibroblast nucleus (Sears, 2004). The heterodimer c-myc/Max is also capable of binding to the DNA E-box motif, competing with the monomer of c-myc for the target genes binding sites (Dang, 1999). The heterodimer c-myc/Max bins to the promoter of *DUSP* gene whose protein dephosphorylates ERK, inhibiting its catalytic activity and closing a negative feedback loop (Chappell et al., 2013) (Figure 1b).

Another target gene of the c-myc/Max heterodimer is the gene *Cdc25A* whose protein product is the phosphatase cdc25A. *Cdc25A* is required for the transition from G_1_ phase to S phase during the cell cycle. In this process, cdc25A removes the inhibitory phosphate groups from threonine and tyrosine residues of G1/S cyclin-dependent kinases CDK4, CDK2 and Cdk1, activating them and allowing cell cycle progression (Figure 1b).

A central problem in the analysis of the codification and transmission of information in this kind of signaling networks *is how the external inputs are correctly coded and decoded to produce specific gene responses* (Díaz and Álvarez-Buylla, 2009). The solution to this question is quit complex because external inputs generate information that flows through the intricate signaling network of target cells. Furthermore, this signaling network (SN) modulates the set of signals that are transmitted to the gene regulatory network (GRN) in a nonlinear form. Both GRN and SN constitute the nonlinear communication channel (CC) of the cell (Zaynab et al, 2015). In this particular case the CC is formed by FGFR, the set of docking proteins Sh2-Sh3, the GAP-Ras circuit, the MAPK cascade, and *C-Myc* with its targets genes. The input to this CC is the extracellular concentration of FGF (Figure 1b).

The FGF CC can be represented with a directed graph formed by *nodes* (genes and proteins that form the CC), and *links* that are the regulatory interactions between the nodes. In a GRN, genes interact with each other by regulatory proteins named transcription factors (TFs), which bind to the promoter sites of target genes to promote or inhibit their transcription. The GRN associated to the FGFR is shown in Figure 1b, in which arrows represent activating interactions and the bars inhibitory interactions. The current state of expression of a gene at time *t* depends on the combination of the activating and inhibitory interactions that act on it. The flow of information between the cell and its environment determines the combination of activated and inhibited genes of a GRN at a particular time *t*, which is the network state in response to a specific stimulus or *input*.

Although a great number of theoretical studies about the dynamics of the MAPK cascade has been published (Kholodenko and Birtwistle, 2009), and the response of ERK to different kind of signals has been tested (Kanodia et al., 2014; Grabowski et al., 2019), there is a lack in our knowledge of how the GRN of the fibroblast activated by the FGFR CC responds to different classes of time-dependent input signals, and how the information flow occurs in each case. Furthermore, cell cycle regulation and cell differentiation show high sensitivity to the amplitude and duration of input signals from growth factors (Kholodenko and Birtwistle, 2009), indicating that gene activation can show a broad range of dynamical responses to time-dependent signals, i.e., the FGFR CC is not only a system of information relay but also a system of information processing used by cells to compute the most accurate gene response to a specific noisy input signal (Díaz and Martínez-Mekler, 2005).

In the present theoretical work, we analyze the response of the GRN formed by the genes *C-Myc, DUSP* and *Cdc25A* to different kinds of FGF input signals (step, quadratic pulses, Dirac delta, and noise) using the tools from the information theory in order to understand how the FGFR CC processes the information from the extracellular medium. To reach this objective, we calculate the Shannon entropy of the emitter and the GRN, and the mutual information between the emitter and the GRN. This analysis also clarifies some clues on how the FGF regulates the cell cycle progression under noisy conditions.

## 2 Model

### 2.1 Stochastic model of a GRN

We define 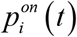 as the probability that the gene *i* of a GRN of *r* interacting genes (*i* = 1, 2,…, *r*) is expressed (state *on*), and 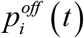 as the probability that the gene *i* is not expressed (state *off*) in response to an input. We assume that the expression or non-expression of the gene *i* under the action of an input *X* is a Bernoulli process and 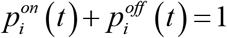.

We propose that the time evolution of 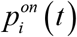 is given in terms of the following master equation (Mousaviana et al, 2016):

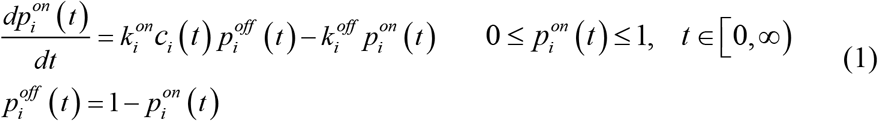

where *c*_*i*_ (*t*) is the concentration of TF molecules in nucleus at time *t, k*_*i*_ ^*on*^ and *k*_*i*_ ^*off*^ are the rate constants for the transition between the expression states.

The Shannon entropy at time *t* due the activation of gene *i* is:

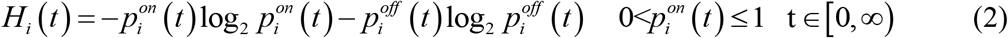

and:

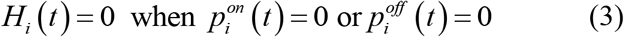

In this form the total Shannon entropy at time *t* due to the current state of activation of the GNR is:

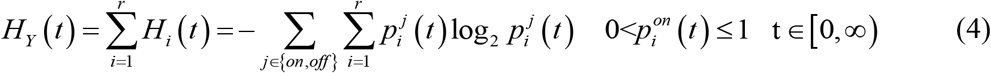

which is the *Shannon entropy* of the receiver *Y*, at time *t*.

In similar way, if *α*(*t*) is the concentration of active receptors at the cell membrane and *α*_*T*_ is the total concentration of receptors, the probability that a receptor molecule is in its active state at time *t* can be approximated for a high number of molecules as:

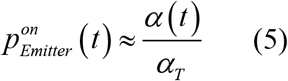

where the total number of receptors is constant at any time.

Thus, the Shannon entropy at the emitter *X* at time *t* is:

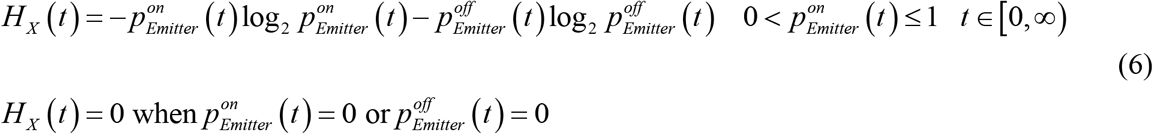

### 2.2 Model of the MAPK cascade

The model presented in this section corresponds to the canonical MAPK cascade without feedback between the cascade components, shown in Figure 1b, and without crosstalk with other components of the SN (Díaz and Martínez-Mekler, 2005).

The first step in the cascade activation occurs as an effect of agonist stimulation with external FGF. The binding of FGF to its receptor form the complex FGF-FGFR, and the cross phosphorylation of two FGF-FGFR forms the dimeric complex (FGF-FGFR)_2_:

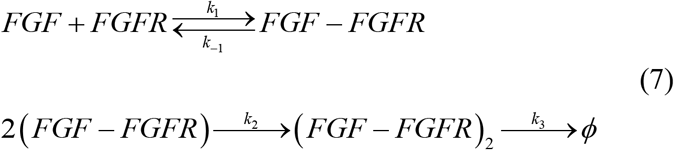

where *ϕ* indicates the internalization of the (FGF-FGFR)_2_ complex for degradation.

Denoting the amount of FGF as *F*, the amount of free FGFR as *R*, the amount of FGF-FGFR complexes as *C*, and the amount of (FGF-FGFR)_2_ complexes as *C*_*2*_ we obtain the differential equations:

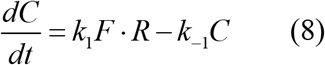

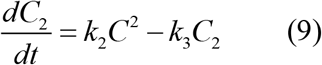

where *k*_*1*_, *k*_*-1*_, *k*_*2*_, and *k*_*3*_ are rate constants (see Table 1),

**Table 1.** Parameters of the model.

| Parameter | Value | Parameter | Value |
| --- | --- | --- | --- |
| $k_1$ | $0.00476 \text{ nM}^{-1}\text{s}^{-1}$ | $SOS_T$ | 90 nM |
| $k_{-1}$ | $0.0008 \text{ s}^{-1}$ | $GAP_T$ | 60 nM |
| $k_2$ | $0.00625 \text{ nM}^{-1}\text{s}^{-1}$ | $ras_T$ | 70 nM |
| $k_3$ | $0.059 \text{ s}^{-1}$ | $raf_T$ | 300 nM |
| $k_4$ | $2 \text{ nMs}^{-1}\text{s}^{-1}$ | $mek_T$ | 300 nM |
| $k_{-4}$ | $0.5 \text{ s}^{-1}$ | $erk_T$ | 300 nM |
| $k_5$ | $10 \text{ nM}^{-1}\text{s}^{-1}$ | $V_{a1}$ | $0.01 \text{ s}^{-1}$ |
| $k_{-5}$ | $1 \text{ s}^{-1}$ | $V_{a2}$ | $0.25 \text{ nM}^{-1}\text{s}^{-1}$ |
| $k_7$ | $0.7 \text{ s}^{-1}$ | $V_{a5}$ | $0.75 \text{ nM}^{-1}\text{s}^{-1}$ |
| $k_{-7}$ | $0.91 \text{ s}^{-1}$ | $V_{a6}$ | $0.75 \text{ nM}^{-1}\text{s}^{-1}$ |
| $k_8$ | $0.025 \text{ s}^{-1}$ | $V_{a9}$ | $0.5 \text{ nM}^{-1}\text{s}^{-1}$ |
| $k_9$ | $0.025 \text{ s}^{-1}$ | $V_{a10}$ | $0.5 \text{ nM}^{-1}\text{s}^{-1}$ |
| $k_{11}$ | $0.025 \text{ s}^{-1}$ | $K_{a2}$ | 8 nM |
| $k_{12}$ | $3 \text{ s}^{-1}$ | $K_{a3}$ | 15 nM |
| $k_{13}$ | $5 \text{ nM}^{-1}\text{s}^{-1}$ | $K_{a4}$ | 15 nM |
| $k_{14}$ | $0.35 \text{ s}^{-1}$ | $K_{a5}$ | 15 nM |
| $k_{15}$ | $1 \text{ s}^{-1}$ | $K_{a6}$ | 15 nM |
| $k_{16}$ | $0.5 \text{ s}^{-1}$ | $K_{a7}$ | 15 nM |
| $k_{17}$ | $0.2 \text{ nM}^{-1}\text{s}^{-1}$ | $K_{a8}$ | 15 nM |
| $k_{-17}$ | $1 \text{ s}^{-1}$ | $K_{a9}$ | 15 nM |
| $k_{18}$ | $0.01 \text{ s}^{-1}$ | $K_{a10}$ | 15 nM |
| $k_{-18}$ | $0.00045 \text{ s}^{-1}$ | $D$ | $1 \text{ s}^{-1}$ |
| $k_{19}$ | $0.005 \text{ s}^{-1}$ | $V_{Cmyc}^{T \max}$ | $8 \text{ nM}^{-1}\text{s}^{-1}$ |
| $k_{-19}$ | $0.001 \text{ s}^{-1}$ | $V_{DUSP}^{T \max}$ | $4 \text{ nM}^{-1}\text{s}^{-1}$ |
| $k_{20}$ | $0.02 \text{ s}^{-1}$ | $V_{CDC25A}^{T \max}$ | $4 \text{ nM}^{-1}\text{s}^{-1}$ |
| $k_{-20}$ | $0.0075 \text{ s}^{-1}$ | $V_{cyt}$ | $800 \mu\text{m}^3$ |
| $k_{21}$ | $1 \text{ s}^{-1}$ | $V_n$ | $400 \mu\text{m}^3$ |
| $k_{22}$ | $0.5 \text{ s}^{-1}$ | $mix_T$ | 5 nM |

In this set of equations, we denote the total amount of FGF applied to the cell at time *t* as *FGFT*. We assume that the total amount of FGFRs (denoted by *R*_*T*_) is a constant, i.e., that the process of FGFR recycling is in a stationary state during the time of simulation. Thus, we can write the mass balance equations:

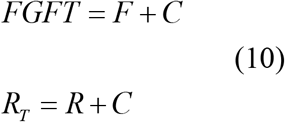

Substituting Eq. (10) in Eq. (8) we finally obtain:

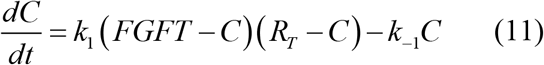

The formation of the (FGF-FGFR)_2_ dimeric complex leads to the activation of SOS through the activation of the docking proteins Shc and Grb2 (see Figure 1). We denote the total amount of SOS as *SOS*_*T*_, the amount of the activated complex Shc-Grb2 as *C*_*Grb2*_, the amount of free SOS as *sos*, and the amount of activated SOS as *sos\**. If we assume that *C*_*Grb2*_ ∝ *C*_2_, we obtain the new equation:

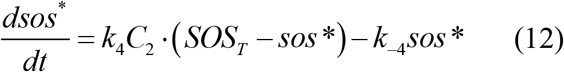

where *k*_*4*_, and *k*_*-4*_ are rate constants (see Table 1).

GAP and Ras proteins form a phosphorylation cycle in which activated SOS promotes the exchange of the GDP bonded to the small G protein Ras for a GTP molecule, while the GAP protein hydrolyses the GTP to GDP. GAP protein is activated by the (FGF-FGFR)_2_ dimeric complex according to the equation:

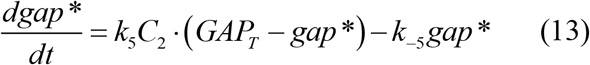

where *gap\** is the amount of activated GAP, *GAP*_*T*_ is the total of GAP in the cell, *k*_*5*_ and *k*_*-5*_ are rate constants (see Table 1).

In this form, the rate of activation of Ras is given by the equation:

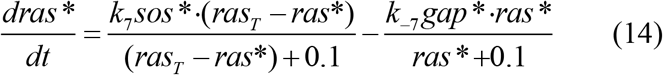

where *ras\** is the amount of activated Ras, *ras*_*T*_ is the total amount of Ras in the cell, *k*_*7*_ and *k*_*-7*_ are rate constants (Klipp et al., 2009)(see Table 1).

Activated Ras targets a MAPKKK that in this case is Raf, the corresponding equation for the rate of activation of Raf is:

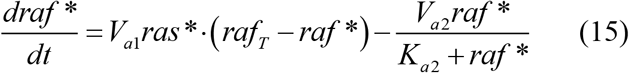

where *raf\** is the amount of phosphorylated Raf, *raf*_*T*_ is the total amount of Raf in cell, *k*_*8*_, *k*_*-8*_ and *V*_*a2*_ are rate constants, and *K*_*a2*_ is also a constant (see Table 1) (Kholodenko, 2000).

Activated Raf initiates the sequence of events of the MAPK cascade that ends with the double phosphorylation of Erk (Kholodenko, 2000) (Figure 1):

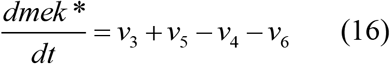

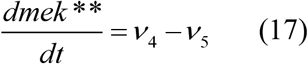

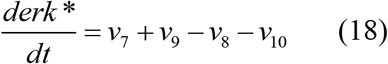

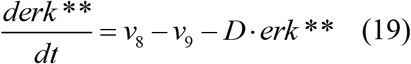

where *mek\** and *mek\*\** are the amount of phosphorylated and double-phosphorylated Mek, respectively. In Eqs. 18 and 19, *erk\** and *erk\*\** are the amount of phosphorylated and double-phosphorylated cytoplasmic Erk, respectively. *D* is the constant of transport of double-phosphorylated Erk into the nucleus.

The variables *v*_*3*_ to *v*_*10*_ in Eqs. 16-19 are the rates of the phosphorylation processes of the kinases in the cascade (Kholodenko, 2000):

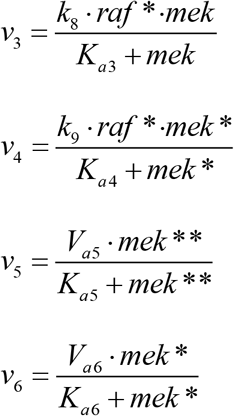

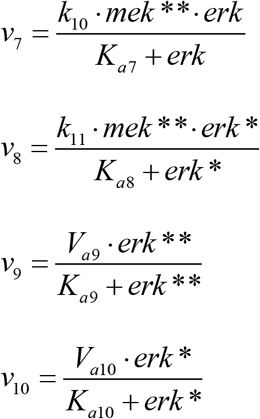

the value of the parameters and rate constants of these equations are the reported in Table 1. Finally, the mass-balance equations for Eqs. 16-19 are:

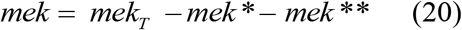

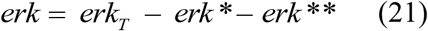

where *mek* is the amount of non phosphorylated Mek, and *mek*_*T*_ is the total amount of Mek in the cell; similarly, *erk* is the amount of non phosphorylated Erk, and *erk*_*T*_ is the total amount of Erk in the cell.

The concentration of Erk in the nucleus must be adjusted because the volume of the nucleus (*V*_*n*_) is lower than the volume of the cytoplasm (*V*_*cyt*_). Hence, the differential equation for the rate of variation of the concentration of Erk in the fibroblast nucleus is:

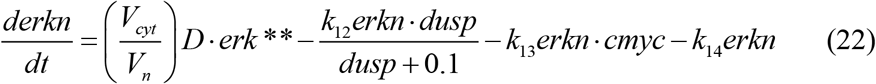

where *erkn* is the amount of doubled phosphorylated Erk in the nucleus, *cmyc* is the amount of protein c-myc, *dusp* is the amount of protein DUSP, *k*_*12*_, *k*_*13*_ and *k*_*14*_ are rate constants (see Table 1). This equation indicates that the rate of variation of the concentration of nuclear Erk is the balance between the rate of transport of phosphorylated Erk from the cytoplasm, minus the rate of interaction of the nuclear Erk with proteins DUSP and c-myc, and the rate of inactivation of nuclear Erk. In this equation, DUSP inhibits the activity of Erk in the nucleus closing a negative feedback circuit (Figure 1).

The rate of variation of the concentration of the protein c-myc in the nucleus is the balance between the rate of translation of the gene *C-Myc* and its rate of degradation:

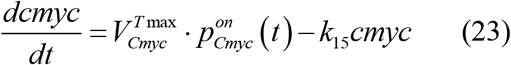

where 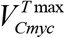 is the maximum rate of translation of the gene *C-Myc*, 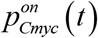 is the probability that the gene *C-Myc* is active at time *t*, and *k*_*15*_ is a rate constant (see Table 1).

The rate of variation of phosphorylated c-myc (*cmyc\**) in the nucleus is the balance between its rate of activation by nuclear Erk, minus its rate of degradation and its rate of interaction with protein Mix:

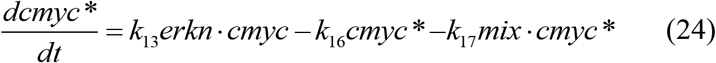

where *mix* is the amount of Mix protein (that we assume is a constant in the model), *k*_*16*_ and *k*_*17*_ are rate constants (see Table 1).

The rate of variation of the concentration of the complex (c-myc)-mix (*ft*) is:

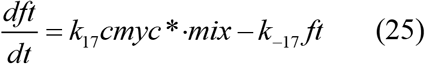

where *k*_*17*_ and *k*_*-17*_ are rate constants (see Table 1).

In this case, the genes *C-Myc, DUSP* and *Cdc25A* form the GRN. The probability of activation of gene *C-Myc* according to Eq. 1 is:

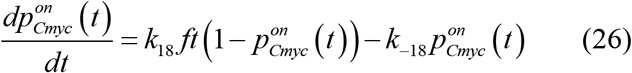

where *k*_*18*_, and *k*_*-18*_ are rate constants.

In a similar manner, the probability of activation of the genes *DUSP* and *Cdc25A* are:

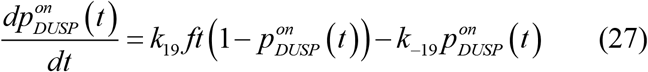

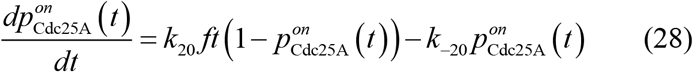

where *k*_*19*_, *k*_*-19*_, *k*_*20*_ and *k*_*-20*_ are rate constants.

The rate of production of protein DUSP (whose concentration is denoted *dusp*) is given by the balance between the rate of translation of the gene *DUSP* minus the rate of inhibition of nuclear Erk by DUSP and its rate of degradation:

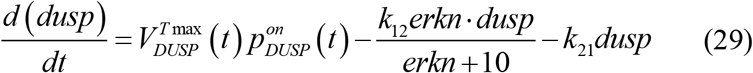

where 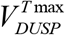 is the maximum rate of translation of the gene *DUSP*, and *k*_*21*_ is a rate constant (see Eq. 16 and Table 1)

Finally, the rate of production of protein cdc25A (whose concentration is denoted *cdc25A*) is given by the balance between the rate of translation of gene *Cdc25A* minus its rate of degradation:

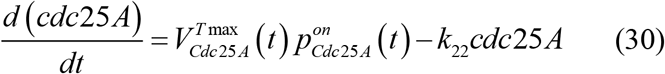

where 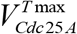 is the maximum rate of translation of the gene *Cdc25A*, and *k*_*22*_ is a rate constant (see Table 1).

The Shannon entropy of the GRN at time *t* can be calculated as:

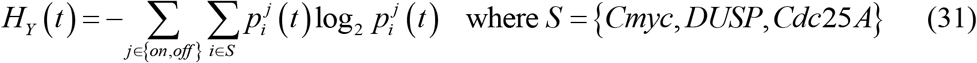

and the total amount of Shannon entropy of the GRN in the time interval (0, τ) of simulation can be calculated as:

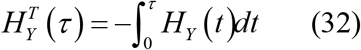

Finally, the Shannon entropy of the emitter at time *t* can be calculated as:

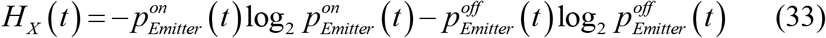

where 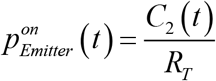. The total amount of Shannon entropy of the emitter in the time interval (0,τ) of simulation can be calculated as:

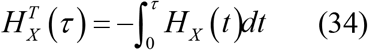

The model was solved using the Euler predictor-corrector method with a time step of 0.05 s. We simulated different time intervals from 120 to 600 minutes to analyze the dynamics of the respond of the cascade subject to different inputs as described below. The initial conditions were set to zero for all the variables of the model, except Raf, Mek y Erk whose initial values were set to 300 nM (Kholodenko, 2000). Protein Mix was held at the constant value of 5 nM during the time of simulation. We used the parameter values of Table 1.

## 3 Results

### 3.1 Dynamics of the FGFR CC response to a FGF step input

We analyzed the dynamics of the CC of Figure 1 in response to a step input signal of 4 nM of amplitude given by the function:

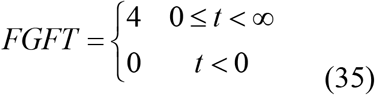

where *FGFT* is the total amount of FGF applied to the cell.

Figure 2a shows that FGF produces a steady level of ∼ 1 nM of the (FGF-FGFR)_2_ dimer, which is the emitter of the signal transmitted by the MAPK cascade. Figure 2b shows in turn that the signal from the emitter produces a steady level of ∼ 6 nM of double phosphorylated Erk in the cytoplasm, a steady concentration of ∼ 8 nM of c-myc in the nucleus, and a peak of nuclear double phosphorylated Erk concentration of 2 minutes of duration with amplitude of 25 nM. This peak is followed by a steady low concentration of nuclear double phosphorylated Erk of ∼ 0.3 nM. Figure 2c shows that all the genes of the GRN have reach a probability of activation higher than 0.9 2 minutes after the signal from the emitter is sent. The probability of activation of the emitter reaches a steady value of 0.3 five minutes after FGF is applied to the cell. The output of the FGFR CC is the temporal distribution of the concentrations of the proteins c-myc, DUSP and cdc25A. Figure 2d show that under FGF saturating conditions the output consist of ∼ 8.5 nM of c-myc, ∼ 7.5 nM of cdc25A and low concentration of ∼ 4.5 nM of DUSP, indicating a probable downregulation of DUSP by double phosphorylated nuclear Erk. Thus, we expect high amounts of proteins c-myc and cdc25A and a low concentration of DUSP in the nucleus.

**Figure 2.**
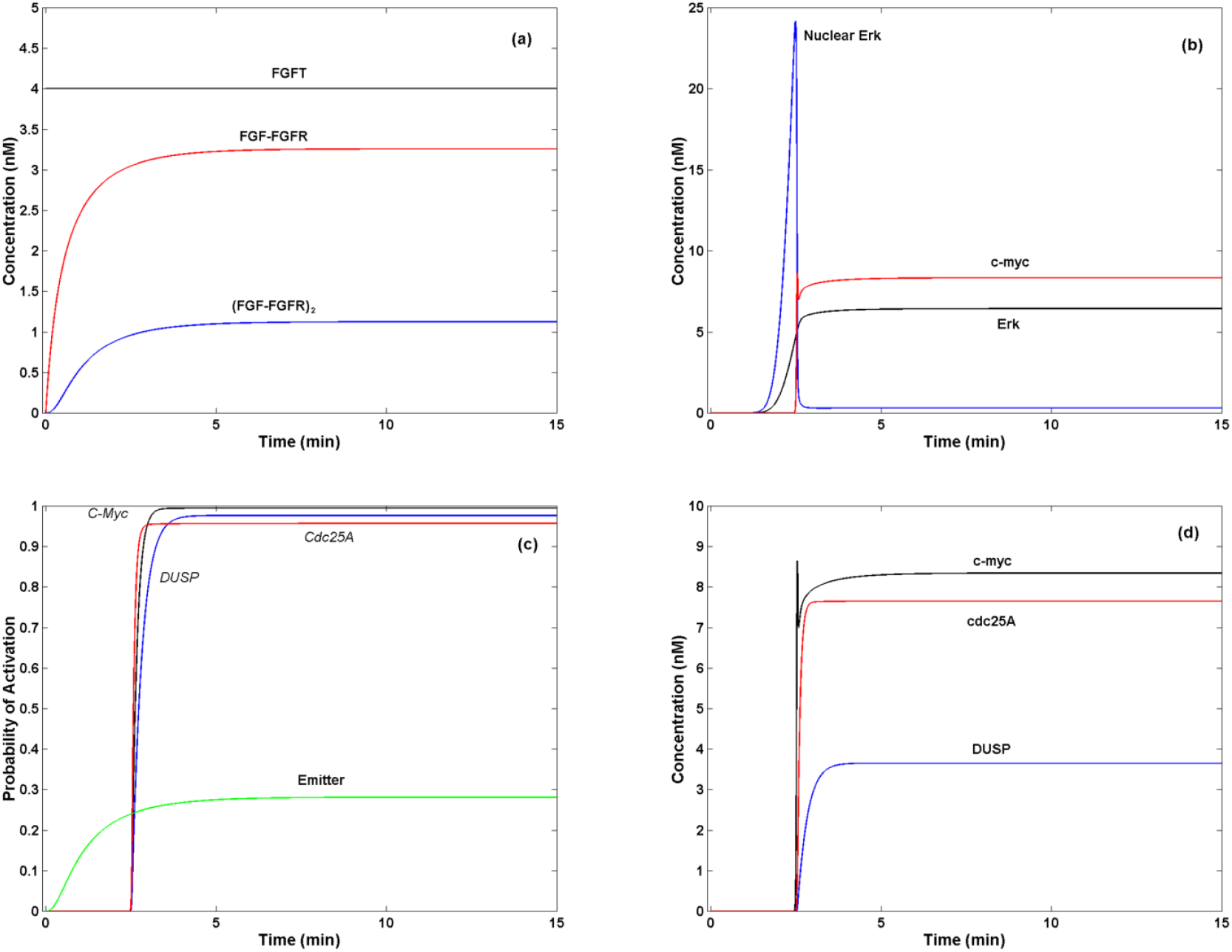
Dynamic of the GRN in response to a FGF step input. (a) 4 nM applied to the extracellular medium during 600 min produces a steady concentration of ∼ 1 nM of the (FGF-FGFR)_2_ complex (emitter). (b) The signal from the emitter is transmitted through the MAPK cascade producing the accumulation of ∼ 6 nM of double phosphorylated Erk in the cytoplasm and ∼ 8 nM of c-myc in the nucleus. Nuclear double phosphorylated Erk initially shows a peak of ∼ 25 nM of concentration that last 2 min until its decay to a steady concentration of ∼ 0.3 nM. (c) The signal from the emitter produces a steady distribution of probabilities in which the three genes of the GRN has a probability of activation of ∼ 1. GRN activation shows a time lag of 2.5 min with respect to the start of the signal at the emitter. In contrast, the probability of activation of the emitter reaches a steady value of ∼ 0.3. (d) The output of the FGF CC in response to the step of 4 nM of FGF consist of a steady distribution of ∼ 7.5 nM of c-myc, ∼ 7.5 nM of cdc25A and ∼ 4 nM of DUSP. We show only the first 15 min of a total of 600 min of simulation.

### 3.2 Flow of information in the FGFR CC in response to a FGF step input

Figure 3a shows that the Shannon entropy of the GRN or *H*_Y_(*t*) has a peak of ∼ 2.75 bits, 2.5 minutes after the signal produced by FGF is emitted and decays to 0.5 bits in ∼ 4.75 minutes. The peak measures the *maximum uncertainty* in the state of activation of the GRN in response to the signal emitted. The curve of *H*_Y_(*t*) follows the temporal course of the variation of the concentration of nuclear double phosphorylated Erk with a slower decay due the presence of an additional low rate process. As shown in Figure 2c, the maximum uncertainty value is reached when the genes of the GRN are not still fully expressed and decay to its minimum value when they are fully activated. In contrast *H*_X_(*t*) smoothly reaches a steady value of ∼ 1 bit five minutes after the activation of the FGF receptor. The total value of *H*_Y_(*t*) (Eq. 26) after 600 minutes is ∼ 1.3 × 10^4^ bits (average bit rate of 21.66 bits/min) while the total value of *H*_X_(*t*) (Eq. 28) is ∼ 3.1 × 10^4^ bits (average bit rate of 51.66 bits/min).

**Figure 3.**
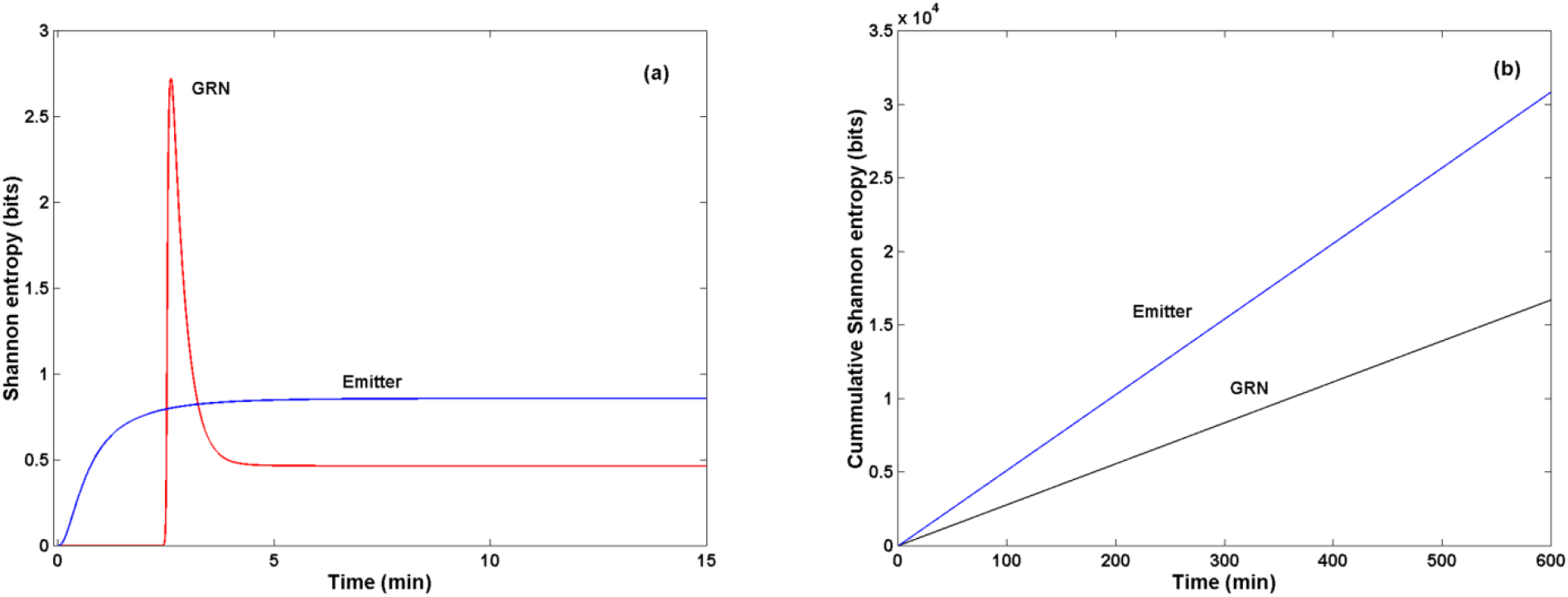
Information flow in the FGFR CC in response to a FGF step input. a) A step **of** 4 nM applied to the extracellular medium during 600 min produces a peak of 2.75 bits of amplitude and 2.5 minutes of duration in the Shannon entropy of the GRN (*H*_Y_(*t*)) after the signal from the emitter arrives to the nucleus. *H*_Y_(*t*) slowly decays to a minimum steady value of 0.5 bits. In contrast, *H*_X_(*t*) reaches a steady value of ∼ 1 bit after the stimulation with FGF. In this set of Figures we only show the first 15 minutes of simulation. b) *H*_*Y*_(*t*) shows an accumulated value of 1.3 × 10^4^ bits and *H*_*X*_(*t*) shows an accumulated value of 3.1 × 10^4^ bits after 600 min of simulation.

### 3.3 Dependence of the FGFR CC dynamics on FGF concentration

We test the effect of the concentration of FGF on the dynamics of the FGFR CC. We found that the CC has not response to *FGFT* < 1.3 nM, showing a threshold value of 1.3 nM, above which the response of the CC increases in intensity as *FGFT* increases in the interval from 1.3 to 1.6 nM, and saturates at 1.6 nM. The response time lag to the stimulation with FGF decreases as the concentration of the agonist increases, reaching its minimum value when *FGFT* = 4 nM. Thus, the full response of the CC is obtained when FGFT = 4 nM.

Figure 4a shows that 1.5 nM of FGFT produces a steady state concentration of ∼ 0.3 of the complex (FGF-FGFR)_2_, which is the emitted signal (Figure 4a). As a result, the concentration of cytoplasmic double phosphorylated Erk and nuclear c-myc shown in Figure 4b are less than the values shown in Figure 2b, in particular the variation of nuclear double phosphorylated Erk is of less amplitude and more duration that the shown in Figure 2b. Thus, the decrease in the concentration of external FGF decreases the intensity of the signal carried by the cytoplasmic double phosphorylated Erk into the nucleus, producing a delay of 10 min in the activation of the GRN and in the time to the full steady activation of the three genes (∼ 15 min) (Figure 4c), which in this case have a low probability of activation: 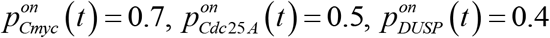. In consequence, the *H*_*Y*_(*t*) curve reaches its maximum value ∼ 10 min post stimulation of the fibroblast cell, and its decaying is slower (Figure 4d) with respect to the curve shown in Figure 3a, reaching a steady value of ∼ 2.3 bits 20 min post stimulation. This result indicates *that the reduction of the concentration of the agonist increments the uncertainty in the state of activation of the GRN in the FGFR CC. H*_*Y*_(*t*) has an accumulated value of ∼ 7.8 × 10^4^ bits after 600 min of simulation (average bit rate of 130 bits/min), which is much higher than in the case of a step of 4 nM of FGF (Figure 3b). As expected, the probability of activation of the emitter is low ∼ 0.1, and *H*_*X*_(*t*) has a lower steady amplitude than in Figure 3b with an accumulated value of ∼ 1 × 10^4^ bits after 600 min of stimulation (average bit rate of 16.6 bits/min)

**Figure 4.**
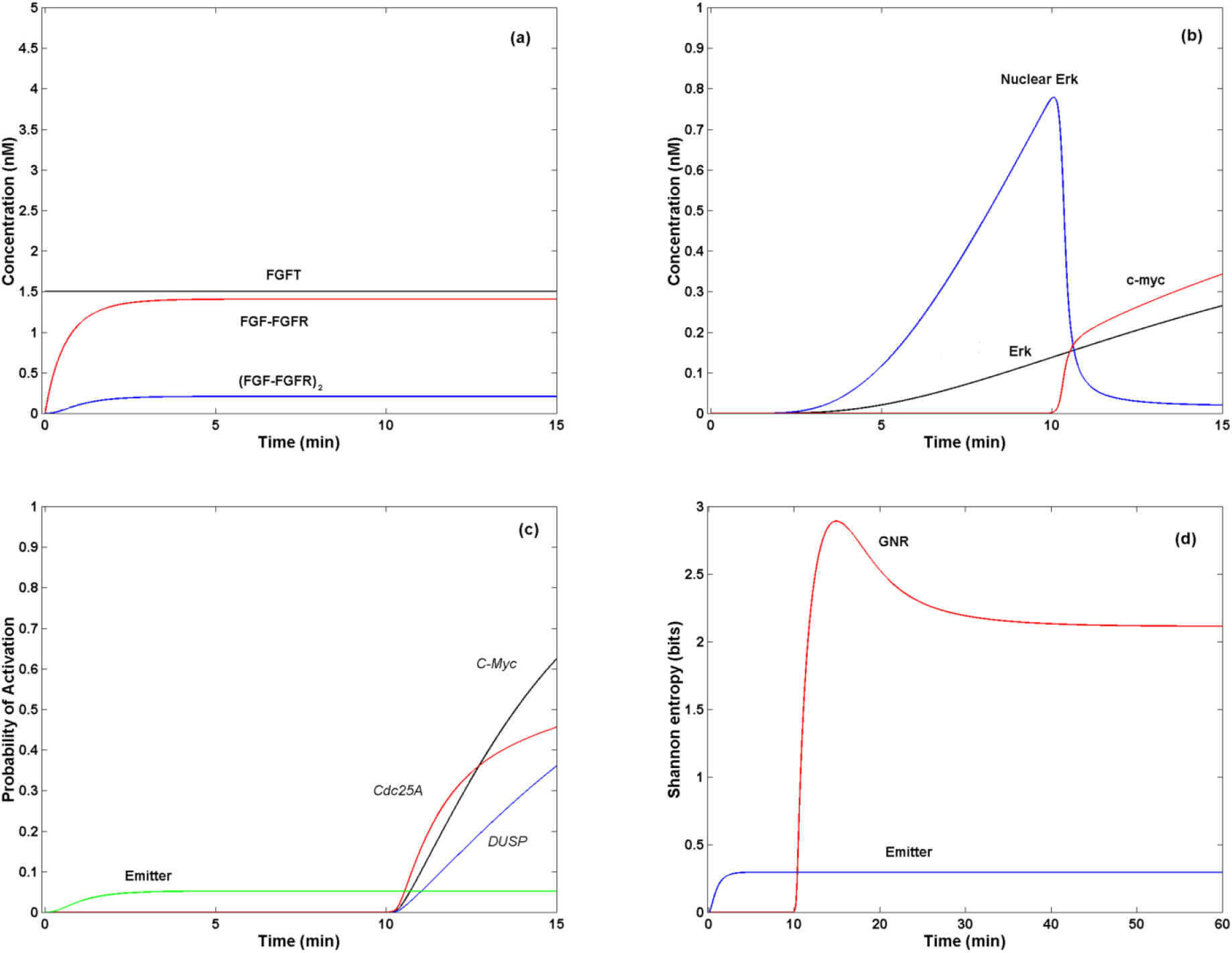
Effect of FGF external concentration on the dynamic of the CC. (a) 1.5 nM of FGF applied to the extracellular medium during 600 min produces a steady concentration of ∼ 0.3 nM of the (FGF-FGFR)_2_ dimer (emitter). (b) The signal from the emitter is transmitted through the MAPK cascade producing the accumulation of ∼ 0.3 nM of double phosphorylated Erk in the cytoplasm and ∼ 4 nM of c-myc in the nucleus. Nuclear double phosphorylated Erk initially shows a peak of ∼ 0.8 nM of concentration that last 12 min until its decay to a steady concentration of ∼ 0.05 nM. (c) The signal from the emitter produces a steady distribution of probabilities in which the gene *DUSP* has a probability of activation of ∼ 0.4, *Cdc25A* has a probability of activation of ∼ 0.5, and *C-Myc* has a probability of activation of ∼ 0.7. GRN activation shows a time lag of 10 min with respect to the start of the signal at the emitter. In contrast, the probability of activation of the emitter reaches a steady value of ∼ 0.1. (d) *H*_*Y*_(*t*) shows a peak of 2.75 bits of amplitude and 10 minutes of duration and slowly decays to a steady value of ∼ 2.3 bits. In contrast, *H*_*X*_(*t*) reaches a steady value of ∼ 0.5 bits after the stimulation with FGF. In this set of Figures, we only show the first 15 minutes of a total of 600 min of simulation.

### 3.4 Response of the GRN to a FGF quadratic pulse

We analyze the dynamics of the GRN subject to a quadratic pulse of FGF defined by the function:

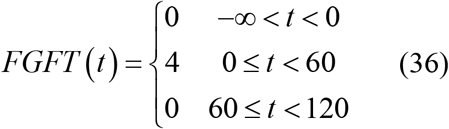

where *FGFT* is measured in nM.

In the time interval *t ∈* (-*∞*, 60) the response of the CC is identical to the shown in Figures 2a – 2c. However, when the signal generated by the emitter is switched *off* in the time interval *t ∈* [60, 120) the emitter, the cytoplasmic and nuclear double phosphorylated Erk, and c-myc are turned *off* in ∼ 25 min (Figures 5a and 5b). The GNR presents a delay in its response because *Cdc25A* turns *off* in ∼ 60 min, *DUSP* turns *off* in ∼ 120 min, and *C-Myc* takes about 200 min to switch *off* (Figure 5c). Unexpectedly, *H*_*Y*_(*t*) shows a peak of ∼ 2.75 bits of amplitude and ∼ 2 min of duration when the CC is turned *on*, and an additional peak of ∼ 2.25 bits of amplitude and ∼ 150 min of duration after the CC is switched *off*. After this second peak *H*_*Y*_(*t*) shows slowly decays to zero when t > 100 min. When the FGF signal becomes zero, the probability of activation of *C-Myc* slowly becomes less than 1, and according to Eq. 31 this gene mainly contributes to the value of *H*_*Y*_(*t*) when the other genes of the GRN shut *off*.

**Figure 5.**
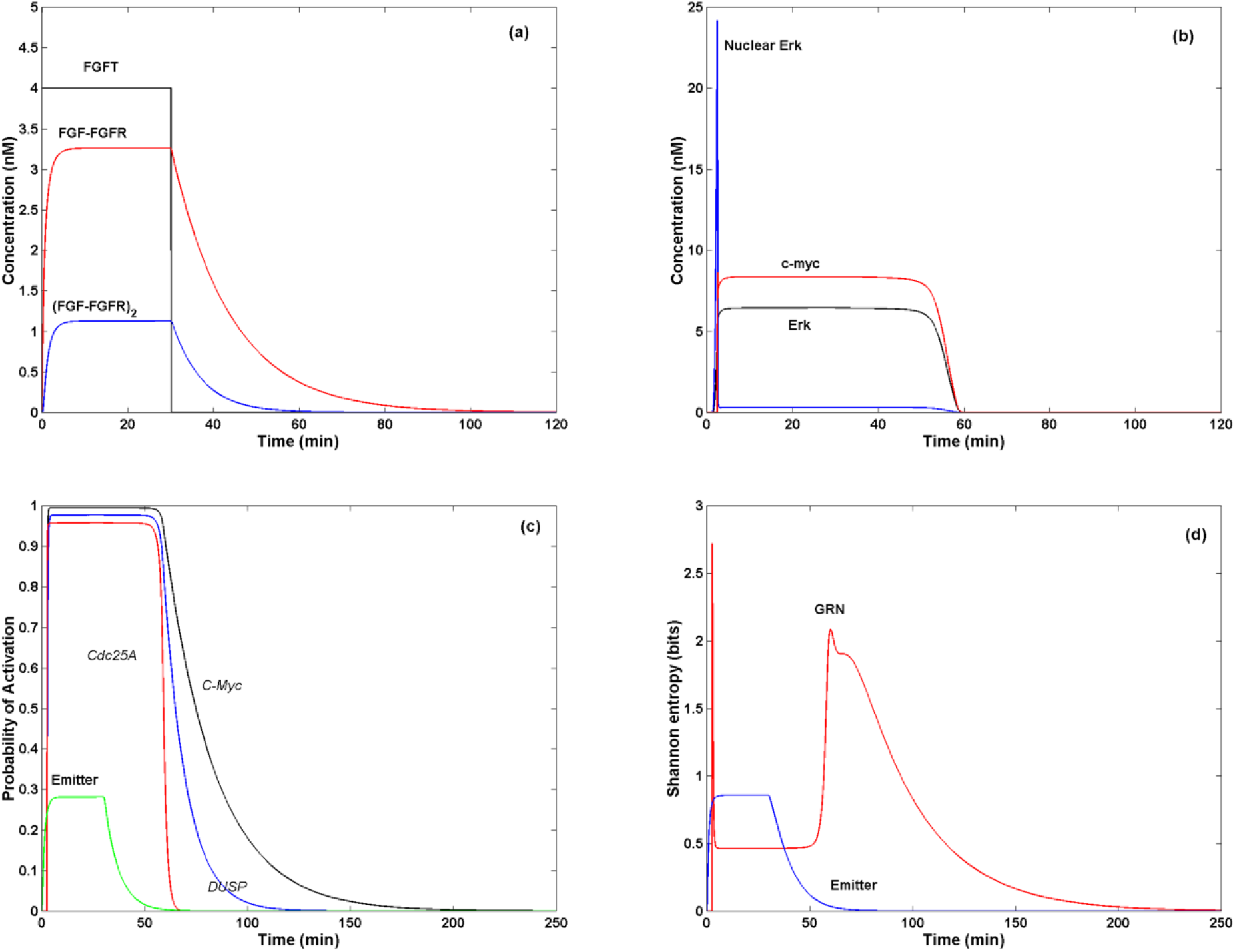
Effect of a quadratic pulse of FGF on the FGFR CC dynamics. (a) FGF applied to the extracellular medium according to Eq. 36 during 600 min produces a quasi-quadratic pulse response of the (FGF-FGFR)_2_ dimer (emitter). (b) The signal from the emitter is transmitted through the MAPK cascade producing a quasi-quadratic pulse response of cytoplasmic double phosphorylated Erk and c-myc, however the dynamics of the nuclear double phosphorylated Erk is not altered. (c) The quadratic input signal from the emitter produces a quadratic response in the probability of activation of the emitter, *DUSP* and *Cdc25A*, but a slow decay in the probability of activation of *C-Myc*. (d) *H*_*Y*_(*t*) shows a peak of ~ 2.75 bits of amplitude and 2 minutes of duration, and a peak of ∼ 2.25 bits of amplitude and ∼ 150 min of duration after the CC is switched *off*. In contrast, *H*_*X*_(*t*) shows a quasi-quadratic response to the input signal. In Figures (a) and (b) we show only 120 minutes of simulation. In Figures (c) and (d) we show 250 min of simulation.

Hence, a quadratic input of FGF produces a quasi-quadratic response in the CC, except for nuclear double phosphorylated Erk and the probability of activation of *C-Myc*. The response of the CC is *quasi-linear* with an output that has a similar form of the input. The presence of an additional peak in the value of *H*_*Y*_(*t*) indicates the possibility of an increase in the amount of uncertainty of the state of the GRN each time the CC is switched *on* or *off*, i.e., *each time that the CC changes its operational state*.

### 3.5 Response of the CC to a train of FGF quadratic pulses

We analyzed the dynamics of the CC subject to a train of quadratic pulses of external FGF with amplitude of 4 nM, period *T* and duration *d* = *T*. In each pulse the CC is switched on during *T*/2 min and then is completely switched off during *T*/2 min.

We found that quadratic pulses of FGF with a period 0 < *T* < 60 min (high frequency pulses) are filtered by the MAPK cascade having not effect on the components of the CC downstream Ras (Figure 6a – 6d). However, quadratic pulses of FGF with a period *T* ≥ 60 min produce a train of quadratic pulses with the same period and a longer duration (Figure 7a – 7d). As *T* → ∞ the response converges to the case of a single quadratic pulse discussed in Section 3.4. In this form, the response of the CC to a train of quadratic pulses is quasi-linear for low frequency pulses, i.e., the MAPK cascade coupled to the GRN behaves like a *low frequency pass filter*. Moreover, the input produces a series of peaks in the value of *H*_*Y*_(*t*) with the same period of the input (Figure 7d). In contrast, *H*_*X*_(*t*) show periodical variations in its value of the same period that the input signal as expected.

**Figure 6.**
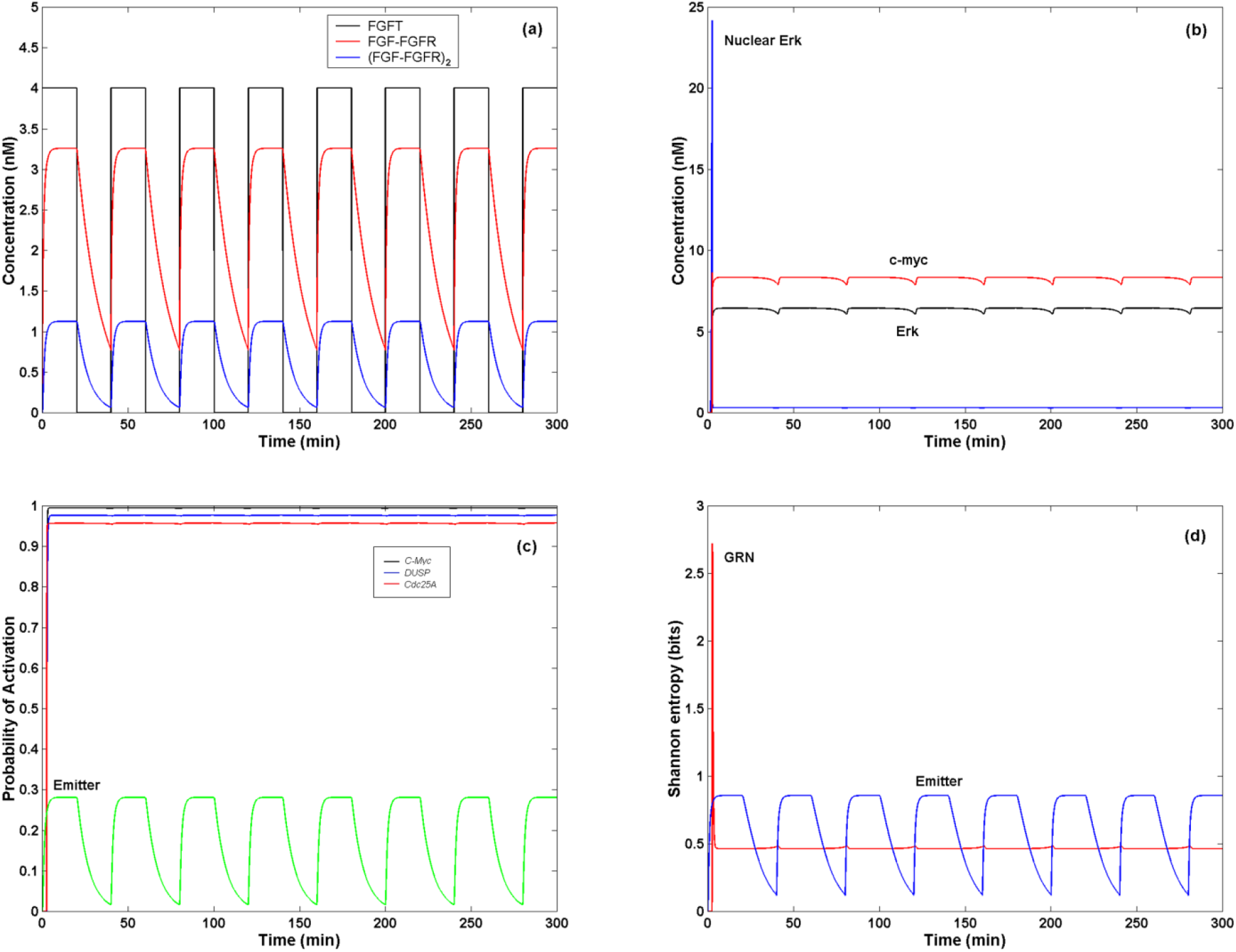
Effect of a train of quadratic pulses with period *T* < 60 min on the CC. (a) Pulses of 4 nM of FGF applied to the extracellular medium every 40 min during 120 min produces a quasi-quadratic pulse periodic response of the (FGF-FGFR)_2_ dimer (emitter), which does not completely turn *off* at the end of each time interval. (b) The signal from the emitter is transmitted through the MAPK cascade and does not produce any affect on cytoplasmic and nuclear double phosphorylated Erk neither on c-myc. (c) The signal from the emitter is transmitted through the MAPK cascade and does not produce any effect on the probability of activation of the genes of the GRN. (d) The signal from the emitter is transmitted through the MAPK cascade and does not produce any affect on *H*_*Y*_(*t*). *H*_*X*_(*t*) shows a quasi-quadratic response to the input signal. In this set of Figures, we show only 300 of the 600 min of simulation.

**Figure 7.**
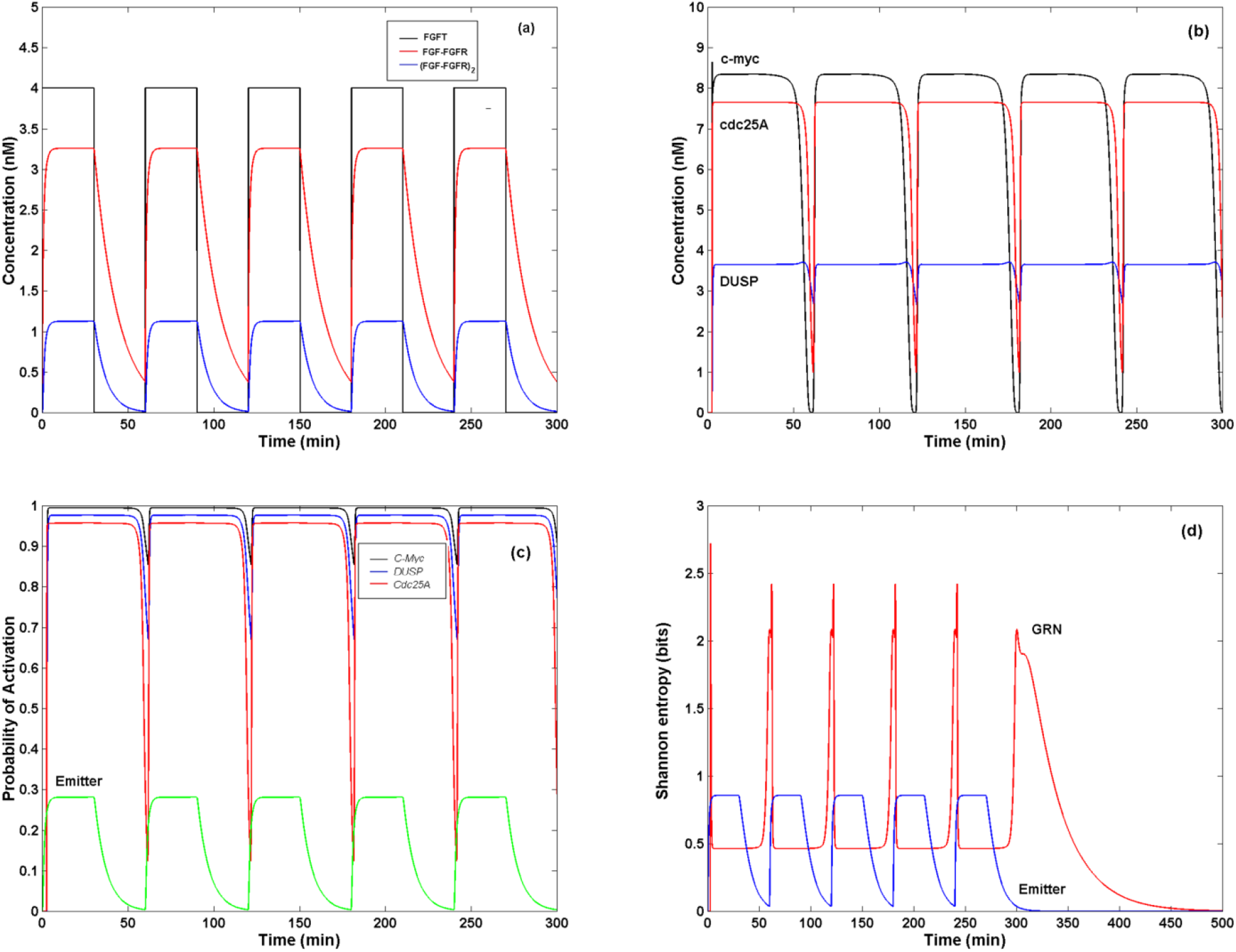
Effect of a train of quadratic pulses of FGF with period *T* ≥ 60 min on the CC. (a) Pulses of 4 nM of FGF applied to the extracellular medium every 60 min during 300 min produces a quasi-quadratic pulse periodic response of the (FGF-FGFR)_2_ dimer (emitter), which completely turns *off* at the end of each time interval. (b) The signal from the emitter is transmitted through the MAPK cascade and produces the periodic variation in the concentration of c-myc, cdc25A, and DUSP proteins, with the same period of the input signal. c) The signal from the emitter is transmitted through the MAPK cascade and produces the periodic variation of the probability of activation of the genes of the GRN, with the same period of the input signal. This Figure shows that the GRN is shut *off* 30 min after the emitter. (d) The signal from the emitter is transmitted through the MAPK cascade and produces a series of peaks in the value of *H*_*Y*_(*t*) with the same period of the input. However, *H*_*Y*_(*t*) reaches the value of zero 200 min after the input has been switched *off. H*_*X*_(*t*) shows a quasi-quadratic response to the input signal. In panels (a) - (c), we show 300 minutes of simulation.

### 3.6 Response of the CC to noise

We found that the FGFR CC acts as a *low frequency pass filter*. Traditionally, experiments and theoretical works (Kholodenko and Birtwistle, 2009; Tka *c̆* ik and Walczak, 2011; Benary et al., 2018) have shown that the ultra sensitivity and all-or-none activation of the MAPK cascade filters noise. However, we have found that when the MAPK cascade is coupled to a GRN this is not completely true because the CC allows the transmission of noisy signals of low rate of variation.

Figures 8a and 8b show that high frequency random signals of FGF has no effect on the steady concentration of the proteins c-myc, cdc25A, and DUSP, In this case, variation of external FGF concentration occur randomly every *τ* minutes, with 0 < *τ* < 50.

**Figure 8.**
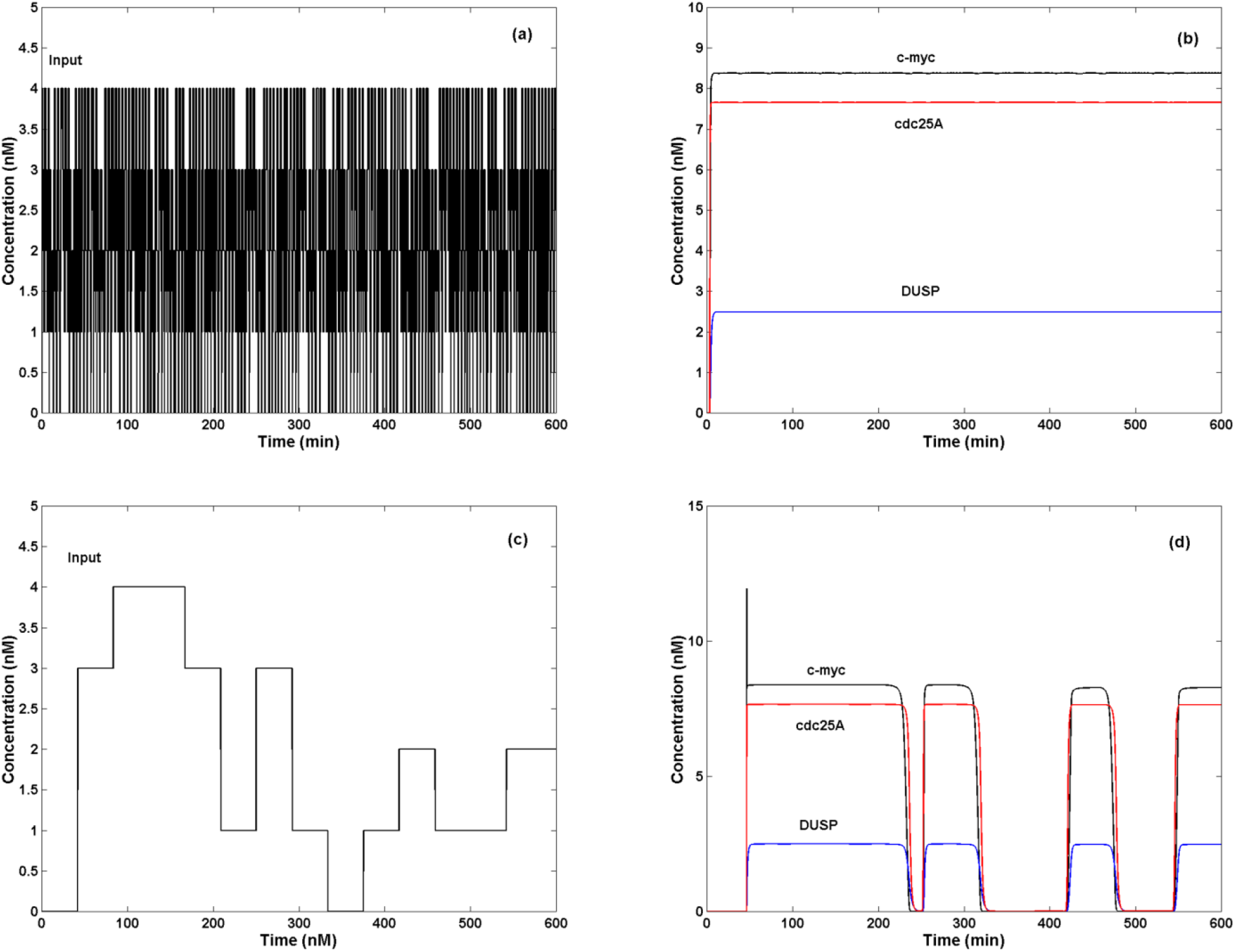
Effect of noise on the CC dynamics. (a) Random signals of high rate of variation of the external concentration of FGF. The signal varies every 0.004 min. (b) Noise of high rate of variation produces a steady concentration of proteins c-myc, cdc25A and DUSP. (c) Random signals of low rate of variation of the concentration of FGF. The signal varies every 50 min. (d) Noise of low rate of variation produces random variations in the concentration of the proteins c-myc, cdc25A and DUSP.

However, Figures 8c and 8d show that a random signal of low rate of variation of the external concentration of FGF produces a random variation of the concentration of the proteins c-myc, cdc25A and DUSP, in intervals of random duration, which is a distorted version of the output shown in Figure 7c. In this case, variation of external FGF concentration occurs randomly every *τ* minutes with *τ* > 50. As expected, the distribution of the probability of activation of the genes of the GRN varies randomly in response to the input signal, given rise to random fluctuations in the value of *H*_*Y*_(*t*) and *H*_*X*_(*t*) (Figure 9). Thus, low rate of variation noise can distort the response of the GNR with undesirable consequences to the cell differentiation process of the fibroblast.

**Figure 9.**
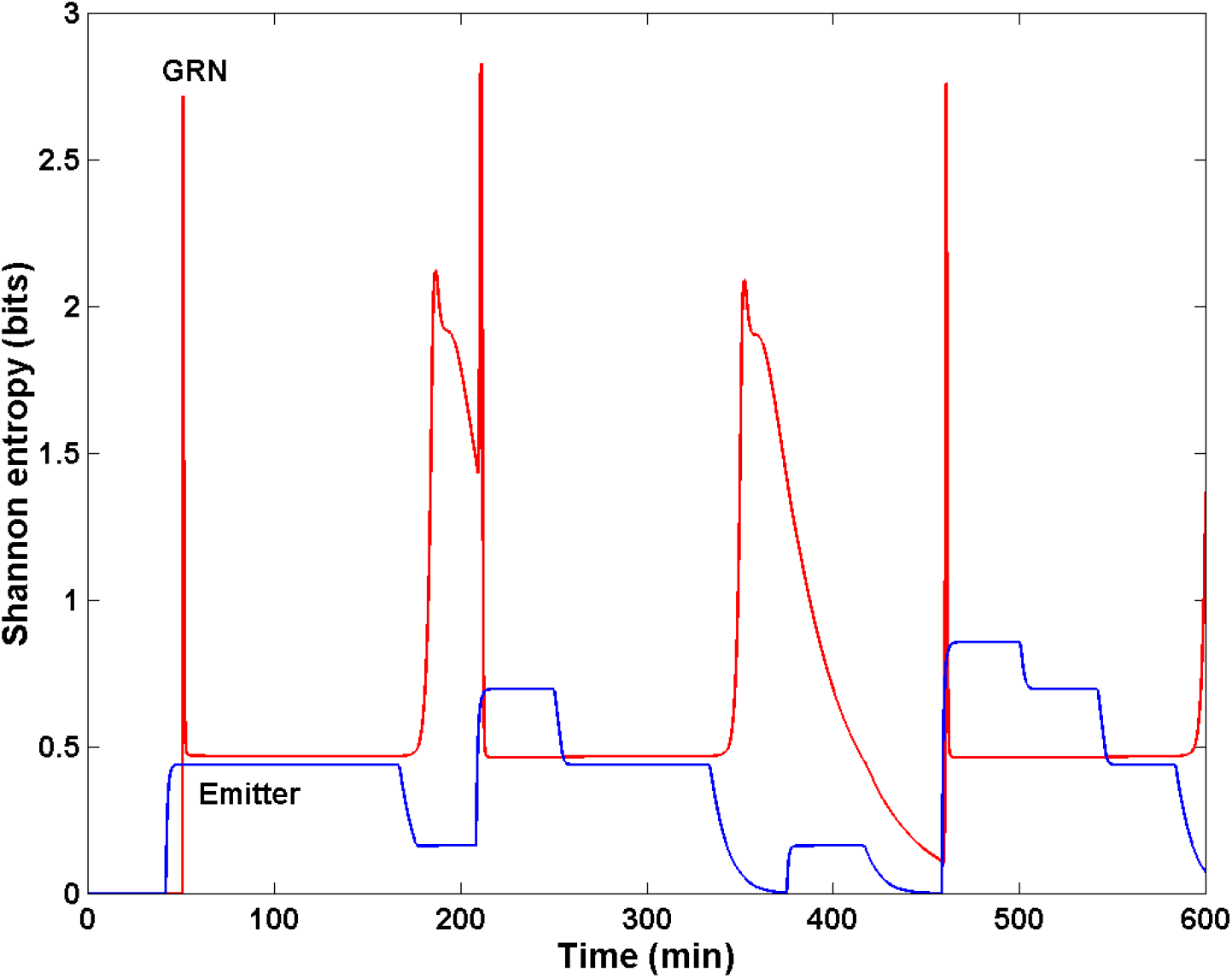
Effect of noise of low rate of variation on the Shannon Entropy of the CC. Random signals of external concentration of FGF that vary every 50 min produces random variations in the value of *H*_*Y*_(*t*) and *H*_*X*_(*t*).

### 3.7 Response of the CC to a Dirac delta

We analyzed the response of the CC to a Dirac delta of FGF given by the function:

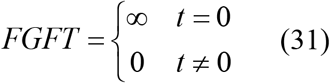

Figure 10a shows the Dirac delta used as input. Figure 10b shows that de response of the CC to this transitory intense signal is the full transitory activation of the GRN giving rise to the transitory production of the proteins c-myc, cdc25A and DUSP during ∼ 40 min.

**Figure 10.**
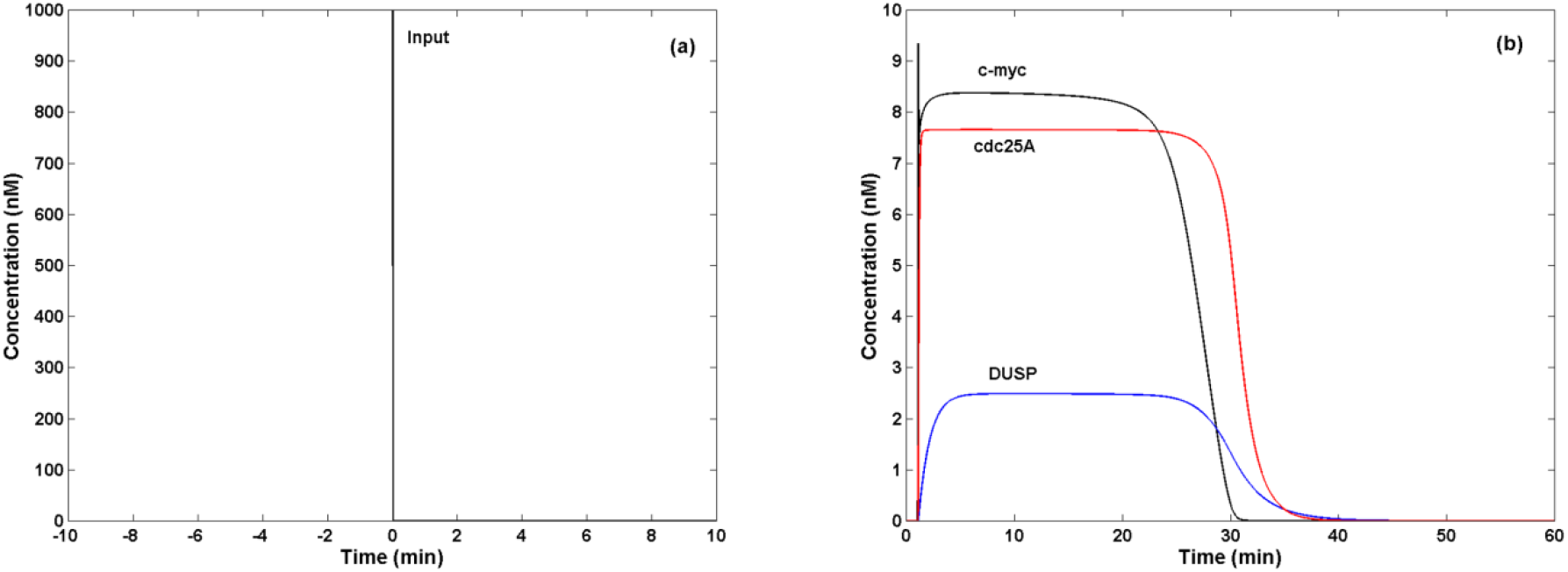
Effect of a Dirac delta of FGF on the CC dynamics. (a) Dirac delta of external FGF. (b) The effect of the Dirac delta (input) is the transitory production of c-myc, cdc25A and DUSP proteins (output) during ∼ 40 min.

## 4 Discussion

Mathematical models in Biology are representations of reality, which implies the simplification of a complex biological process into a set of central components, which we assume are the key determinants of the spatio-temporal dynamics of the system under study (Eftimie, et al., 2016). This set of main components form a molecular network, in which the number of nodes and links, together with the distribution of links between nodes, determine the network dynamics (Eftimie, et al., 2016; Díaz, 2020). In the particular case of the coupling of a SN with a GRN, the resultant network can be considered as a molecular CC. The main function of this kind of CC is to mediate in the transmission of information from the cell surface to the GRN, and generate a specific genetic response. Because GNRs are control systems, the analysis of the flow of information in this type of CCs deals with a problem of control and communication (Bi and Noel, 2020; Wiener, 1961).

In this sense, beside the extensive research that has clarified the main features of a great number of manmade CCs, there are features of the biological CCs that are just beginning to be understood. For example, in the FGFR CC the coding of information is made using chemical reactions (Tka *c̆* ik and Walczak, 2011). Thus, the input signal is the temporal variation of the external FGF concentration, which is coded in the number of (FGF-FGFR)_2_ complexes formed for each concentration of external FGF (Figures 3a and 4a). Hence, the emitter of the original signal is the (FGF-FGFR)_2_ dimer, which is coupled to the MAPK or transmitter, and the information is coded in the number of activated dimers at time *t*.

Although the MAPK cascade interacts with the SN, and the cascade itself has feedback circuits between its components, we decided not to take into consideration these interactions in our model in order to determine the function of this basic molecular structure of the cascade in the modulation of the signal transmitted by it. Previous works (Kholodenko, and Birtwistle, 2009), have shown that the MAPK cascade has ultrasensitivity and filtering properties that reduce the effects of noise on the signal transmitted by the cascade, increasing the fidelity of the transmission. Thus, the practically all-or-none activation of the cascade avoids the distortion of the transmitted signal by cytoplasmic noise (Tka *c̆* ik and Walczak, 2011; Vaseghi, 2000). We found that the MAPK cascade also filters external noisy signals of high rate of random variation (Figures 8a and 8b). However, we unexpectedly found that the cascade allows the transmission of periodic signals of low frequency (Figure 7) or signals with low rate of random variation (Figures 8c and 8d), acting like a *low frequency pass filter*.

In the FGFR CC the receiver of the signal is the *C-Myc* GRN, which is activated by the double phosphorylated Erk that enters the nucleus (Figures 1 and 3b). In this form, the signal is translated into a specific duration of the activated state of the target genes *C-Myc, DUSP* and *Cdc25A*. Thus, we proposed a stochastic model of the process of gene transcription instead of a deterministic one (Eqs. 26-28), which we consider a better approach to the form in which gene translation occurs in real GRNs. In consequence, the input signal is *decoded* into a specific time-dependent probability distribution of gene activation for each concentration of the emitter:

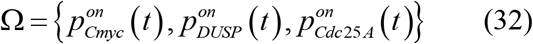

that give rise to a temporal distribution of their respective protein concentrations (Eqs. 23, 29 and 30). These proteins are the *effectors* of the information generated by the input. Summing up: *in the FGFR CC each external concentration of FGF is coded into the concentration of the (FGF-FGFR)*_*2*_ *dimer, which activates the cascade in an all-or-non manner, transmitting the information from the dimmer to the nucleus where is decoded into a probability distribution of activation of the genes of the GRN that finally gives rise to a specific distribution of concentration of the effector proteins* (Díaz and Alvarez-Buylla, 2009; Mousaviana et al., 2016).

A fundamental question arises at this point: is the specific distribution of concentration of these proteins the correct output for each input signal? The stochastic nature of gene expression produces a certain level of uncertainty about the real state of activation of the GRN at time *t* that we can measure with the Shannon entropy *H*_*Y*_(*t*) (Eq. 31). This quantity measures the contribution of the lost of information in the real state of activation of each gene of the GRN at time *t* with regard to the state of activation of the overall GNR in response to a given input. This loss of information or uncertainty is due to noise and other molecular processes in which the genes take part. Thus, in our model *we will never know the exact state of activation of a GNR*, and we propose that the output reflects the *most probable* specific distribution of concentration of the proteins c-myc, DUSP, and cdc25A in response to a particular input. This flow of information between the emitter and the GRN to produce a specific response is measured with the mutual information *I* _(*X*;*Y*)_(*t*) between *X* and *Y*.

In order to know the mutual information between the emitter and the GNR in the CC, we assume in Eq. (31) that:

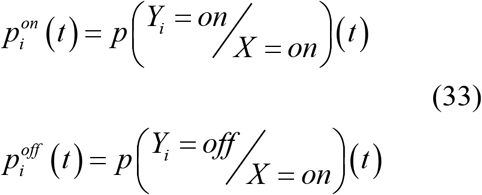

where the random variables *Y*_*i*_ = state of activation of gene *i* at time *t, i* ∈{*C*-*Myc, DUSP,Cdc*25*A*} and *X* = state of activation of the emitter at time *t*, can take only the values *on* and *off*.

When FGF is applied to the CC, then the emitter is in state *on*, and:

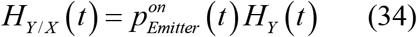

We calculate the mutual information between the random variables *X* and *Y* under FGF stimulation as:

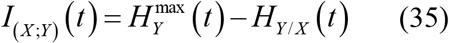

where the value of 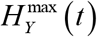 is estimated from the output of the model.

We obtain from Eq. (33):

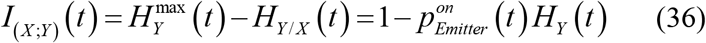

The capacity of a CC measures the ability of the receiver to discriminate between two different signals and produce the correct output in a noisy channel. Formally, the channel capacity is the maximum rate at which information can be sent with arbitrarily small probability of error. The Epidermal Growth Factor Receptor (EGFR) CC has a capacity of 1.84 bits/h (Benary et al., 2017; Grabowski et al., 2019). We determinate the FGFR CC capacity *χ* using the result from Eq. (36) in the following equation:

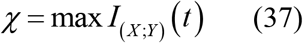

Figure 11a shows the graphic of *I*_(*X;Y*)_(*t*) when a *step* of 4 nM of FGF is applied to the CC. The maximum value of *I*_(*X;Y*)_(*t*) is ∼ 2.9 bits during 3 min, thus the capacity of the CC (Eq. (37) is estimated in a value of ∼ 0.96 bits/min. It is possible that an increase in the complexity of the GRN coupled to the MAPK cascade enlarges the capacity of the FGFR CC in this case.

**Figure 11.**
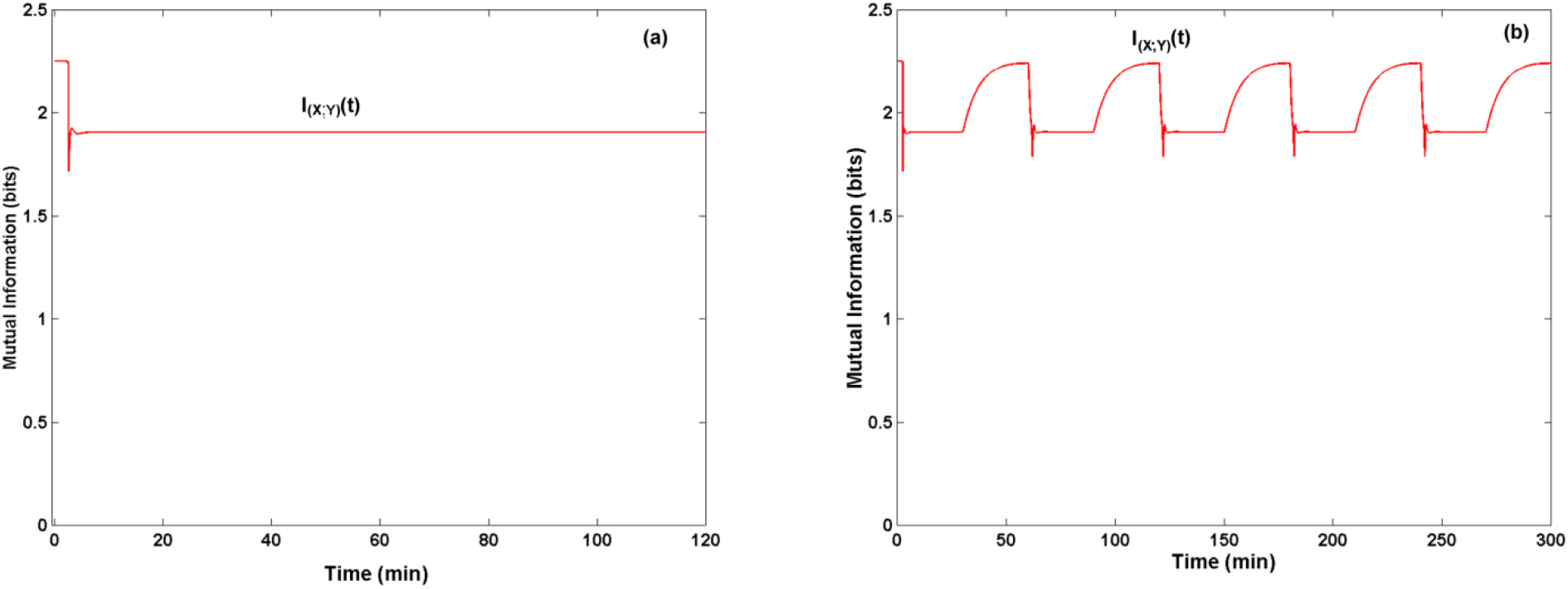
Mutual information of the FGFR CC. (a) Mutual information graphics of the FGFR CC in response to step of 4 nM of FGF. The figure shows a maximum value of ∼ 2.9 bits, a minimum value of ∼ 2.25 bits and a steady value of ∼ 2.75 bits. (b) Mutual information graphics of the FGFR CC in response to a train of quadratic pulses. The figure shows that in this case the mutual information oscillates between a maximum value of ∼ 2.9 bits and a minimum value of ∼ 2.3 bits with a periodicity of *T* = 60 min.

Our results show that c-myc regulates cell cycle progression of the fibroblast (Bretones et al., 2014) down regulating the nuclear concentration of the dual phosphatase DUSP and enhancing the nuclear concentration of the phosphatase cdc25A (Figure 2d). Although DUSP can dephosphorylate the threonine/serine and tyrosine residues of nuclear Erk, down regulating its kinase activity, Erk can also phosphorylate DUSP and down regulate its activity. For the set of parameter values of Table 1, *our model predicts the down regulation of DUSP protein activity in the nucleus, and a decrease in the probability of activation of the DUSP gene (Figures 2, 3 and 4), when the fibroblast is stimulated with FGF*.

In contrast, the enhanced concentration of the phosphatase cdc25A and the high probability of activation of its gene *Cdc25A* (Figures 2c and 2d) induced by c-myc indicates that the stimulation of the fibroblast with FGF favours its proliferation. Protein cdc25A acts over the complex cyclin D-cdk4 and cyclin E-cdk2 allowing the G1/S phase transition, and also regulates G2/M transition before its degradation (Lim and Kaldis, 2013; Shen and Huang, 2012). In this form, the model predicts that *any step signal of FGF above the threshold concentration of ∼ 1*.*3 nM induces cell cycle progression* from phase G1 to phase S, and from phase G2 to M, when DUSP is down regulated by double phosphorylated nuclear Erk (Figure 1).

The above scenario is supported by the results shown in Figure 5, in which a quadratic pulse of FGF induces the transitory production of cdc25A and DUSP proteins that possibly stops the transition from F1 to S due to the inactivation of the *Cdc25A* gene after 100 min. In this second scenario, *an intense transitory stimulation of the fibroblast with a quadratic pulse of low duration of FGF above the threshold value of ∼ 1*.*3 nM is not enough to overcome cell arrest in G1 phase*. Experimental results indicate that cdc25A protein has a half life of ∼ 20 min, thus it must be produced continuously during at least ∼ 10 hrs to ensure G1/S transition and during ∼ 20 hrs to allow the G2/M transition (Shen and Huang, 2012; Timofeev et al., 2009). Additionally, an intense stimulation of the fibroblast with a Dirac delta of FGF produces a transitory response of the GRN of ∼ 40 min of duration (Figure 10), which is insufficient to overcome the fibroblast arrest in the G1 phase.

Therefore, the low frequency pass filtering properties of the FGFR CC ensures the continuity of the cell cycle progression avoiding the effects of periodic regular variations of high frequency (T < 60 min) (Figure 6) or noise of high rate of random variation (Figures 8a and 8b) on the concentration of c-myc and cdc25A. However, periodic regular variations of low frequency (Figure 7) or noise of low rate of random variation (Figures 8b and 8c) could obstruct cell cycle progression by continuously switching *off* cdc25A production.

The filtering property of the FGFR CC can be an important feature to avoid intense and high frequency interferences on the signals carried by the MAPK cascade to the nucleus ensuring the fidelity of the information transmitted to the GRN from the (FGFR-FGFR)_2_ dimer complex. However, we are using a simplified model of the FGFR CC and possibly there are complementary molecular mechanisms that damp the effect of low frequency interferences on cell cycle progression, but the presence of this type of response of the CC to this type of signals is an unexpected result that *opens the possibility of a real mechanism that regulates cell proliferation*.

During G1/S progression the emitter and the GNR have a steady mutual information of ∼ 2.75 bits (Figure 11a), i.e., the amount of information that the state of activation of the GRN gives about the state of activation of the (FGFR-FGFR)_2_ dimer is ∼ 2.75 bits, value that is constant during all the process and measures the mutual nonlinear dependence between the random variables *X* e *Y*. Thus, once the GNR is activated with a step of 4 nM of FGF the CC reaches a steady state with a constant level of uncertainty about the correct transmission and interpretation of the message elicited by the FGF at the cell surface, indicating that while the fibroblast stimulation with a step of FGF continues the G1/S transition does not require of additional interchange of information between the emitter and GRN to complete the cell cycle progression to phase S.

This scenario changes when the CC is subject to a train of quadratic pulses of 4 nM of FGF with a period of *T* = 60 min. Figure 11b shows a periodic variation in the value of *I*_(*X,Y*)_(t) from 2.4 to 2.9 bits during the interval of time that the CC is switching *off*, while the peaks in the value of *H*_*Y*_(t) occur just at the moment in which the CC turns *off* (Figure 7d). This result indicates that there is a transitory transference of information between the GRN and the emitter, which probably is the gain in the amount of information about the state of activation of the emitter obtained from the GRN during its turning *off*, these peaks of information probably coincide with the intervals of time in which the G1/S transition is interrupted (Figures 7 and 11b). Figures 8d and 9 clearly show that random noise of low rate of variation completely distorts the flow of information in the CC probably producing the arrest of the cell cycle or an irregular progression of the transition from G1 to S.

## 5 Conclusions

Cell cycle progression is the central process in cell proliferation and must be rigorously controlled to allow the correct fibroblast tissue replacement, its regeneration after an injury and to avoid cancer. In this process, extracellular FGF triggers the activation of the genes responsible of this control by the activation of the intracellular signaling network that carries on the signal generated by the (FGF-FGFR)2 dimmer, or emitter, to the nucleus. An essential component of this signaling network is the MAPK cascade, which in our work is the transmitter of the signal from the cell surface to the nucleus and closes the FGFR communication channel. The MAPK cascade has properties of ultrasensitivity that traditionally is related to its function as noise filter that ensures a high fidelity of the information transmitted by it. However, most of the works that sustain this affirmation are based on theoretical and experimental models in which the cascade is uncoupled to a GRN. In our work we analyze the flow of information in the FGFR CC when the signals are transmitted by the MAPK cascade to the *C*-*Myc*-*DUSP*-*Cdc25A* GRN, in absence of crosstalk with other components of the signaling network, and we found that low frequency noise (period *T* ≥ 60 min) is not filtered and can modify the output of the CC, i.e., the observed amount of the effector proteins c-myc, cdc25A and DUSP is continuously distorted by the noise. An additional effect is that low frequency noise can stop cell cycle progression from G1 phase to S phase because the required threshold value concentration of cdc25A is not continuously sustained in the nucleus during the 10 hours that the G1/S transition last. In consequence, our model suggest that the sustained activation of the FGFR CC with a step of FGF > 1.3 nM, or with periodic pulses of FGF with a period T < 60 min, is required for cell cycle progression. During the G1/S transition the amount of uncertainty at the GRN in response to a step of 4 nM of FGF remains at a steady value of ∼ 2.75 bits, value that increases every time that the state of the GRN varies, indicating an additional interchange of information between emitter and GRN to complete the cell cycle progression to S phase. From our model, we could make a rough estimation of the capacity of the FGFR CC in 0.96 bits/min, value that we think probably changes with the size of the GNR.

## 6 Author Contributions

Both authors contributed equally to this work.

## 8 Acknowledgments

JD thanks Erika Juarez Luna for logistical support.

**References:** Kholodenko (2000), Klip et al. (2009), Huang and Ferrell (1996).

